# A molecular representation to identify isofunctional molecules

**DOI:** 10.1101/2024.05.03.592355

**Authors:** Philippe Pinel, Gwenn Guichaoua, Nicolas Devaux, Yann Gaston-Mathé, Brice Hoffmann, Véronique Stoven

## Abstract

The challenges of drug discovery from hit identification to clinical development sometimes involve addressing scaffold hopping issues, in order to optimize biological activity or ADME properties, improve selectivity or mitigate toxicology concerns of a drug candidate, not to mention intellectual property reasons. Docking is usually viewed as the method of choice for identification of isofunctional molecules, i.e. highly dissimilar molecules that share common binding modes with a protein target. However, in cases where the protein structure has low resolution or is unknown, docking may not be suitable. In such cases, ligand-based approaches offer promise but are often inadequate to handle large-step scaffold hopping, because they usually rely on the molecular structure. Therefore, we propose the Interaction Fingerprints Profile (IFPP), a molecular representation that captures molecules binding modes based on docking experiments against a panel of diverse high-quality protein structures. Evaluation on the Large-Hops (*LH*) benchmark demonstrates the utility of IFPP for identification of isofunctional molecules. Nevertheless, computation of IFPPs is expensive, which limits the scalability for screening very large molecular libraries. We propose to overcome this limitation by leveraging Metric Learning approaches, allowing fast estimation of molecules’ IFPP similarities, thus providing an efficient pre-screening strategy applicable to very large molecular libraries. Overall, our results suggest that IFPP provides an interesting and complementary tool alongside existing methods, in order to address challenging scaffold hopping problems effectively in drug discovery.

## 1 Introduction

### Context

The journey from hit discovery to a clinical drug is loaded with various challenges, including the occasional need to address scaffold hopping problems. ^1^ While hit molecules may have been identified for example in through High Throughput Screening, they do not always constitute viable drug candidates as they might suffer from poor ADME properties. Adverse toxicology profiles or poor selectivity can also appear later in the development process. Different degrees of scaffold hopping have been characterised by ^2^ to depict how much the searched molecule needs to differ from the starting point. Small-to medium-step scaffold hopping can be handled by trained medicinal chemists or conventional chemoinformatic approaches such as QSAR. They usually imply keeping key substituent (-R) groups that interact with the targeted protein, while changing the connecting fragments using classical tricks such as swapping of carbons and heteroatoms in heterocycles, heterocycles ring opening or closure, leading to a new molecule that retains some similarity with the hit compound. In large-step scaffold hopping, the new molecule needs to share very limited structure similarity with the original hit, for example when the hit presents many flaws, or when its overall scaffold is protected by a patent, thus restricting future commercial exploitation. The goal is then to roam the chemical space in search for new molecules that interact with the protein target according to binding modes that are similar to those of the initial hit. Pairs of molecules that meet this criterion can be called isofunctional molecules (as a synonym of large-step scaffold hopping pairs of molecules), referring to the fact that they share a similar mechanism of action with respect to a protein target. Identification of isofunctional molecules is a challenging problem that usually requires the use of computational methods. This definition excludes for example pairs of enzyme inhibitors, one binding to the active site and the other to an allosteric site, or one being reversible and the other irreversible (forming a covalent bond withing the active site). Indeed, there would be no rationale that could be exploited to discover the second molecule of such pair when starting from the first.

When the 3D structure of the protein target is known, docking (a structure-based method) is the standard approach used to solve these cases, and has proved successful in the literature. ^3,4^ However, when the resolution of the 3D structures is low, or when only apo structures are available, docking approaches may suffer from uncertain predicted poses, and consequently, from low confident docking score. Furthermore, the 3D structure of the target may be unknown. Various approaches are available to predict the overall 3D models for proteins, including the efficient AlphaFold algorithm. ^5^ However, these models may not be reliable at the level of structural details such as the orientation of side-chain or backbone loops, although these details are critical for the performance of docking approaches. ^6^

In the present article, we tackle the most difficult setting, i.e. identification of isofunctional molecules when the structure of the target protein is unknown, and where novel dedicated computational methods are the most critically needed. This particular setting leaves room for ligand-based approaches. These methods use as input molecular representations that are characterized by descriptors such as ECFP, MACCSkeys, FTrees, or 2D and 3D pharmacophores. These representations proved to help identify new hits. ^7–10^ Overall, they rely on the principle that similar molecules are likely to bind to the same protein. However, most classical molecular descriptors are not adapted to solving large-step scaffold hopping problems, because they tightly depend on the chemical structure of molecules, while isofunctional molecules should lie in remote regions of the chemical space.

Therefore, the problem of finding isofunctional molecules boils down to defining an encoding that maps molecules into a space where molecules that bind to the same pocket are close to each other. This encoding needs to be as much as possible agnostic of the 2D structure of molecules, in order to allow molecules that are dissimilar in terms of chemical structure to be close in the feature space. In this context, molecular representations based on biological properties are expected to be better suited to large-step scaffold hopping. Several encodings have been described in the literature. ^11,12^ Some, like CBFP, ^13^ encode predicted activities for a profile of assays, thus defining bioactivity fingerprints for molecules. However, many of these bioactivity fingerprints need to be predicted for most molecules, because the corresponding assays were conducted on a limited number of molecules, and the corresponding prediction models may lack generalization properties. Consistent with this remark, pre-training a convolution neural network that predicts protein-ligand interactions based on the PCBA dataset that contains a profile of 90 bioactivities for thousands of molecules, did not improve the prediction performances of the algorithm. ^14^

### Contributions

In the present paper, we propose a novel biological representation of molecules inspired from such profiles that can be used as input by ligand-based methods in order to solve large-step scaffold hopping problems. This encoding integrates information about protein-ligand interactions according to docking experiments against a panel of proteins for which a 3D structure of high quality is available. We refer to this molecular profile as the *Interaction Fingerprints Profile* (IFPP) in the following. We evaluate the interest of this representation in addressing large-step scaffold hopping problems using the Large-Hops (*LH*) benchmark, ^15^ as detailed below. Finally,to address the computational cost associated with this representation, we employ Deep Metric Learning, which allows very fast pre-screening of large molecular libraries.

## 2 Results

### 2.1 Using the *LH* benchmark to assess the interest of IFPP as molecular representation to identify isofunctional molecules

To assess the relevance of the IFPP molecular representation for addressing scaffold hopping compared to other reference methods, we used the *LH* benchmark ^15^ (https://github.com/iktos/scaffold hopping) as a tool to compare different classical molecular representations. This database comprises 144 pairs of ligands for 69 diverse proteins. These pairs are known examples of isofunctional molecules that bind similarly to the same protein while displaying highly dissimilar 2D structures. For each ligand pair, 499 decoys were carefully picked to avoid bias towards either of the ligands. In particular they possess similar global physical and chemical properties than the ligands, while they are as distant from each ligand of the pair as these ligands are from each other, in terms of chemical structure.

With the *LH* benchmark, the performance of computational methods are evaluated as follows: for each of the 144 pairs, one ligand is designated as the known active while the other is considered as the unknown active, and added to a pool of 499 decoys. Given the known active, each method ranks the unknown active among the 499 decoys, the lower the rank, the better. For each pair, one molecule or the other can be considered as the known active, which leads to 288 scaffold hopping cases to solve. The performances are compared according to the proportion of successful cases, i.e. for which the unknown active is retrieved in the top 5%, and to their cumulative distribution of ranks for the unknown active.

Our proposal involves comparing molecular representations based on the performance of their associated similarity measures. More precisely, for each case in the *LH* benchmark, the unknown active will be ranked among the decoys based on its similarity to the known active. In cases of tied ranks, we assigned the average of the ranks. Similarity measures will be compared based on their proportion of successful cases. The similarity measures leading to the best success rates will correspond to the molecular encodings that are best adapted to solving scaffold hopping problems.

However, it is important to note that in real-life applications, where various active and inactive molecules could be available for the target of interest (which is not the case in the *LH* benchmark), we would recommend searching for isofunctional molecules using the most promising encodings as input of more sophisticated ligand-based computational methods such as QSAR or machine learning models.

### 2.2 The Interaction Fingerprints Profile for molecular representation

In this section, we describe the motivation for the proposed IFPP as molecular representation and explain how it is computed. We then evaluate its relevance for solving scaffold hopping cases with ligand-based approaches, using the *LH* benchmark.

#### 2.2.1 Rationale

As recently depicted in, ^15^ none of the classical ligand-based methods relying on structural features of molecules display good performances in the challenging task of solving large-step scaffold hopping problems.

We suggest that molecular features derived from interactions that can be formed between a given molecule and protein pockets may be more relevant to solving this problem. Indeed, by definition, isofunctional molecules display dissimilar structures but share similar binding modes with a targeted protein. When the 3D structure of the target is available, docking can be used to search for candidates sharing similar binding modes. Our assumption is that even when the 3D structure of the target is unknown, the tendency to form similar interactions could be observed in other proteins.

The proposed approach is related to ”ensemble methods” in machine learning, and particularly to ”weak learners” methods. ^16^ A weak learner is a model that performs only slightly better than the random prediction for a given task. In other words, it captures limited signal about the task at hand. Alone, it is useless, especially when compared to a ”strong learner” that captures much of the signal, thus achieving good accuracy on the considered task. Unfortunately, such strong learner is often either too hard to train, or even inaccessible. However, when aggregating several weak learners that may be easier to access, the performances of the resulting ensemble model can reach those of the strong learner. As an example, this general principle corresponds to the theory behind random forest algorithms. ^17^ With this concept in mind, we introduce the following analogy. The strong learner would be docking in the protein target: based on the hit molecule binding mode, docking can be used to search for highly dissimilar molecules that present similar binding modes with this protein pocket. This strong learner is unavailable for a protein of unknown 3D structure. In such cases, the weak learners consist in docking the molecules in other proteins of known 3D structure. More precisely, we assume that two isofunctional molecules for a given protein would present a tendency to form similar interactions with proteins, in general, and that this tendency could be detected by docking. Docking a molecule in various proteins would allow to detect interactions between this molecule and protein pockets. Using ”enough” weak learner, i.e. docking in ”enough” proteins could be used to define the Interaction Fingerprint Profile of the molecule. The IFPP could then be used in ligand-based methods, replacing the unavailable strong learner (i.e. docking in the protein of interest).

It is important to note that the docking experiments used to build the IFPPs can be viewed as pure simulations. We are primarily interested in understanding how molecules would interact with the considered pockets, rather than whether they are true ligands with high affinity for these proteins.

In summary, our proposal relies on the idea that, within the space defined by IFPPs, two isofunctional molecules would be closer to each other than to randomly chosen decoys. They would also be closer to each other in this space than in the space defined by chemical descriptors.

#### 2.2.2 Building the IFPP of molecules

##### Principle of IFPP computation

The IFPP is built from weak learners that correspond to docking molecules into a panel of proteins of known 3D structures. The number and the nature of the proteins in this panel must be defined. We assume that a more diverse protein panel will cover a wider range of potential interactions that molecules can form.

We used the DOCK6 software ^18^ to dock molecules in a panel of selected protein pockets. For each pocket, the best scored pose is retained (see more in Section 4.3), and the interactions are retrieved using a proteinligand interactions detection algorithm, as detailed in ^15^ and in Section 4.2. The considered interactions are: hydrogen bond, weak hydrogen bond, halogen bond, salt bridge, multipolar, hydrophobic, pi-hydrophobic, pi-amide, pi-cation and pi-stacking. We derive an Interaction Fingerprint for the molecule in the considered pocket, in the form of a fixed-size binary vector that incorporates, for each residue in the binding site, the types of interactions it forms with the molecule. The final IFPP of the molecule is defined by concatenating these binary Interaction Fingerprints for the proteins belonging to the panel. Figure 1 illustrates how the IFPP of a molecule is obtained.

**Figure 1:**
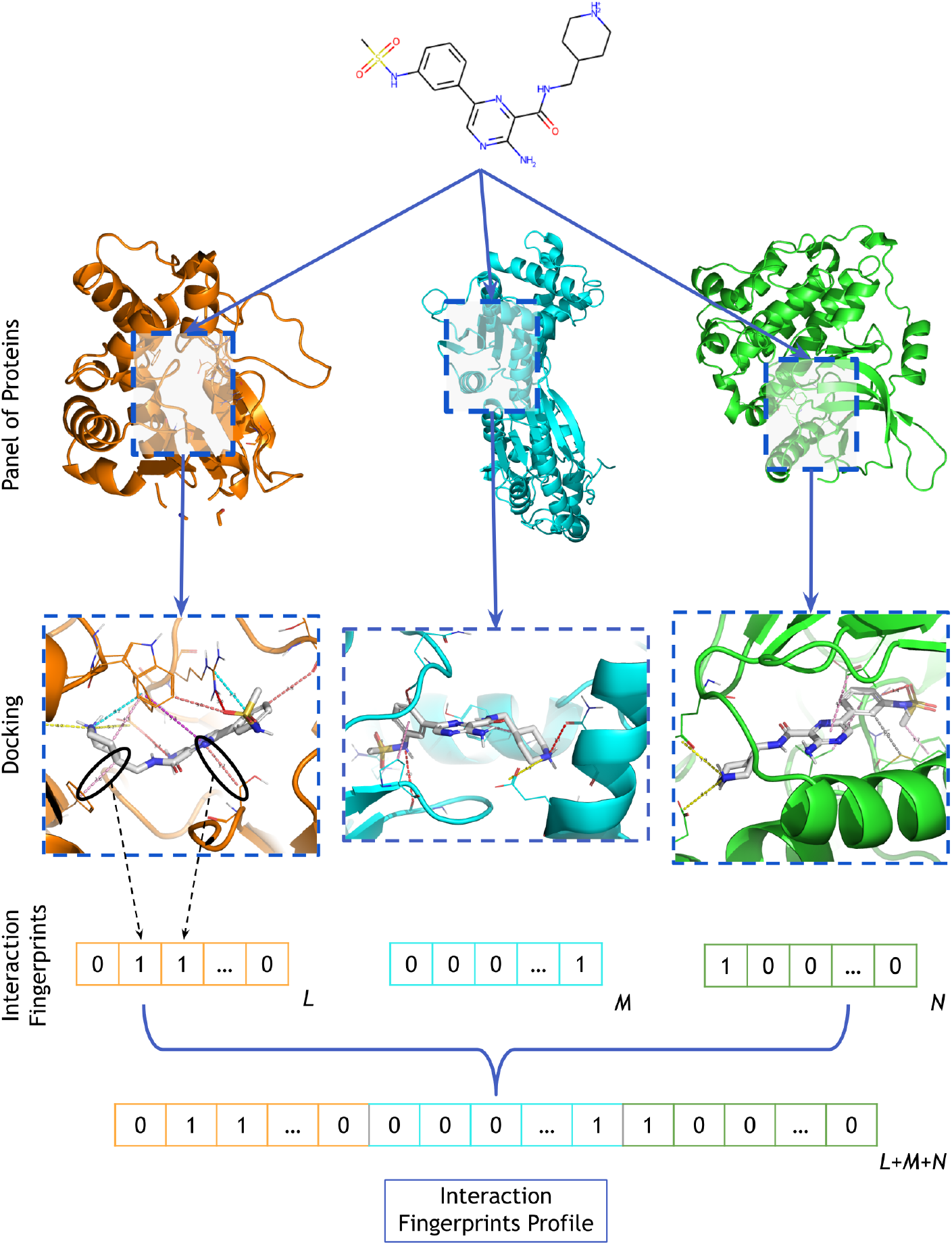
Illustration of how molecular IFPPs are built. In this example, a panel of 3 proteins is chosen. First, the molecule is docked in each of the proteins. Then, interactions of the best scoring pose are retrieved and encoded in a binary interaction fingerprint. *L, M* and *N* represent the length of the interaction fingerprints, dependent of the size of the binding site. Finally, interaction fingerprints are concatenated to form the final IFPP of length *L* + *M* + *N* .

##### Defining a ”minimal” size for the panel of proteins used to compute the IFPP

As detailed in Section 4.1 of Materials and Methods, we selected a panel of 69 proteins that span all superfamilies of protein structures, according to the SCOP database. ^19^ These proteins were selected from the PDBbind database, ^20^ using successive filters. In particular, they are known to be ”druggable”, with available 3D structures in complex with ”drug-like” ligands. In addition, we only kept structures for which the calibration of the docking protocol was achieved, according to a successful re-docking of the bound ligands. None of those proteins is present in the *LH* benchmark.

However, real-life studies may require screening millions of molecules, and docking large chemical libraries against 69 proteins to derive the IFPPs would lead to high computational costs. Therefore, we undertook a preliminary study on the *LH* benchmark. The goal was to assess whether we could reduce the size of the IFPP (i.e. consider a smaller number of proteins to build the IFPP) without degrading the performance of its associated similarity measure, as introduced in Section 2.1. The corresponding similarity measure between molecules was defined as the Tanimoto similarity of their IFPPs.

This preliminary study would require 34, 569 docking experiments for each scaffold hopping pair (2 actives and 499 decoys docked in 69 pockets), and consequently, almost 5 million docking experiments for the whole *LH* benchmark. To reduce these computational costs, for each scaffold hopping case, we kept 49 randomly picked decoys (out of 499), among which the unknown active was ranked according to its similarity with respect to the known active. We considered that a successful experiment corresponded to ranking the unknown active in the top 5% molecules (out of 49 + 1 = 50 molecules, thus corresponding to a rank ≤ 2.5). We tested different sizes of protein sets from the initial panel, ranging from 10 to 65. We performed 100 random draws of pockets for each set size from the 69 initially selected proteins.

As shown in Figure 2, the performance of the IFPP increases with the size of the protein set. We do not seem to reach a plateau yet, suggesting that considering a higher number of proteins would improve the performances. However, as a compromise between computational cost of the IFPPs and performances of the associated similarity measure, we kept a panel of 37 proteins randomly picked in the 69 initially selected protein. The list of these proteins is provided in Supplementary Material.

**Figure 2:**
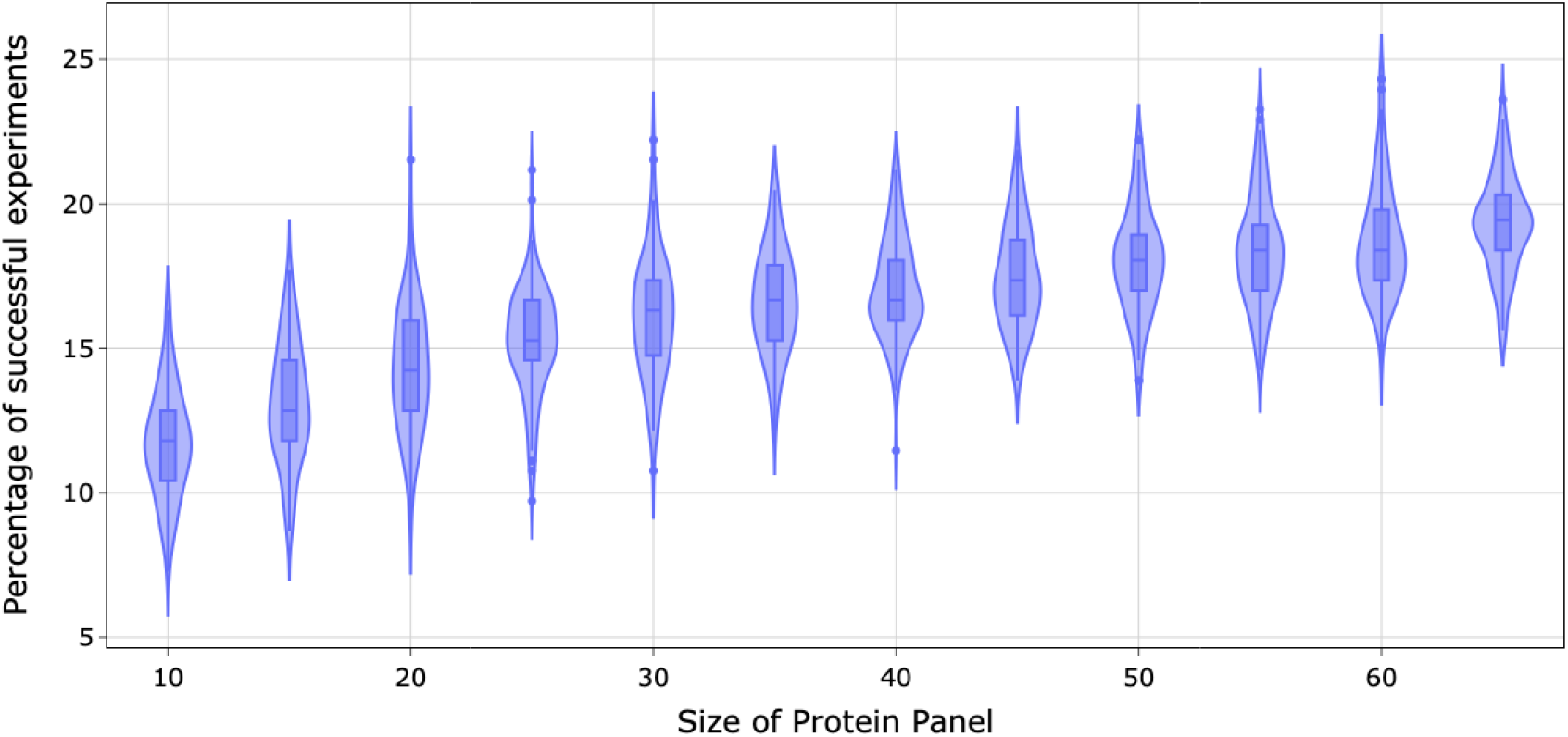
Influence of the size of the protein panel used to define the IFPP on the performance of the associated similarity measure on a reduced *LH* benchmark. Successful experiments are defined by a rank below 2.5 for the unknown active.

#### 2.2.3 Performance of IFPP on *LH* benchmark

As introduced in Section 2.1, one method to evaluate the effectiveness of molecular representations for solving scaffold hopping cases is by comparing the performance of their associated similarity measures in the *LH* benchmark.

In addition to the IFPP, we considered several classical structure-related encodings: Morgan fingerprints and 2D Pharmacophore fingerprints computed with RDKit, ^21^ as well as 3D Pharmacophore and 3D Shape computed with Pharao. ^22^

Figure 3 shows the performances of the similarity-based methods using these 5 encodings. All methods display modest performances on this benchmark, with success rates below 25% for all methods, which illustrates that solving large-step scaffold hopping cases is a difficult task. As expected, 3D methods tend to perform better than 2D methods, because ligand binding is a process that occurs in the 3D space. Interestingly, the similarity in IFPP outperforms the others.

**Figure 3:**
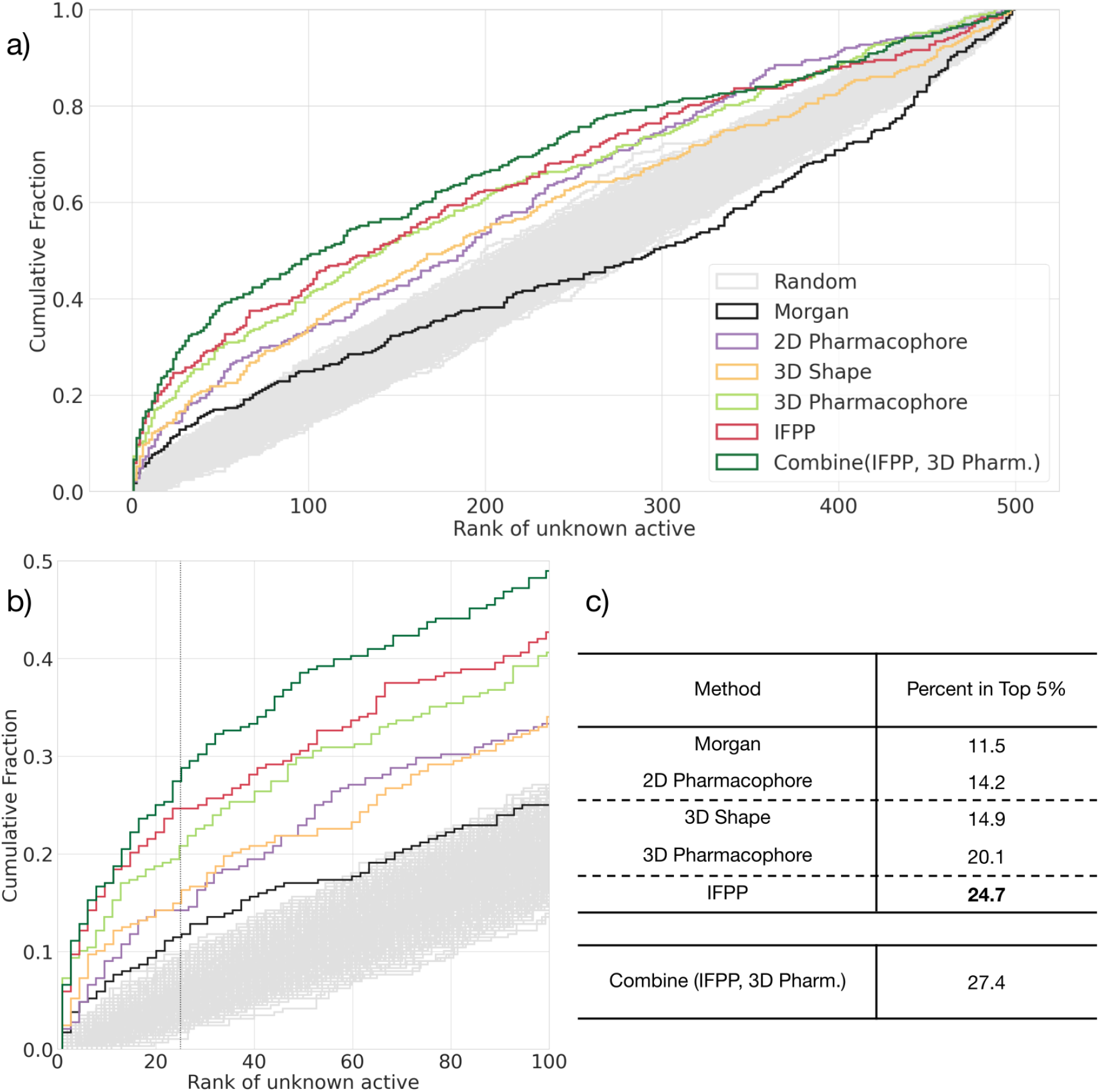
Results on the *LH* benchmark. The cumulative histogram curves of each similarity-based method are plotted in a). A zoom of the same graphs is provided in b) with vertical grey lines corresponding to ranks of top 5% ranks. Table c) displays the percentage of successful scaffold hopping problems for similarity-based methods using various molecular representations, according to a rank of the unknown active in the top 5%.

We performed the Kolmogorov-Smirnov test with the alternative hypothesis being that the cumulative distribution of ranks of the IFPP is greater than that of the 3D Pharmacophore. This resulted in a p-value of 0.038, a limited but significant improvement.

We also computed the Spearman correlation of ranks between all methods, and summarised the results in Table 1. It shows that the IFPP is uncorrelated to the 3D Pharmacophore, and also to other methods. This indicates that this new representation captures information about the protein-ligand interactions that is absent in the other considered molecular representations. Because of their low correlation, we can combine the IFPP and 3D Pharmacophore representations, according to the minimum rank from both methods. This allowed to improve the performances, as illustrated in Figure 3 with a success rate of 27.4% in the top 5%.

**Table 1:**
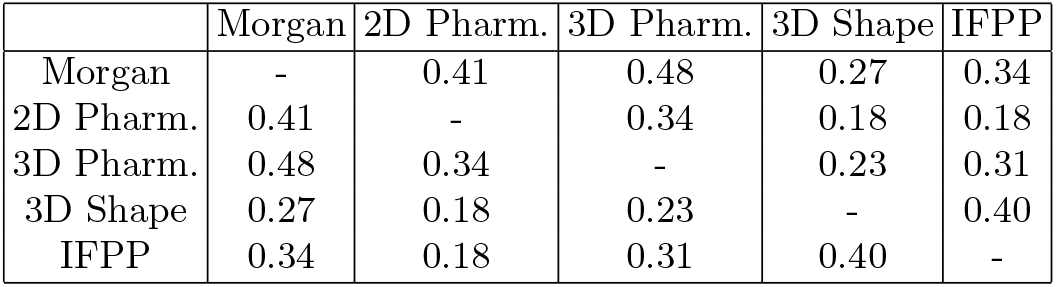
Spearman correlations between the rank of the unknown active for the similarity-based methods using various considered molecular representations.

We confirmed that the IFPP similarity was not successful only in specific families of proteins, by computing the mean, standard deviation and quartiles ranks it assigns to the unknown active for scaffold hopping cases involving proteins from different superfamilies. A superfamily, as defined by the SCOP database, ^19^ gathers proteins with different sequences, but that may have a common evolutionary origin according to their structures and functional features. For example, in the case of the kinase superfamily, in 75% of the 94 related scaffold hopping cases (out of 288 cases), the unknown active was ranked above 36.8, i.e. not in the top 5% best ranked molecules.

However, a drawback of the IFPP is its computational cost. Indeed, building the IFPP of a molecule requires its docking in 37 protein pockets. For example, to perform our study on the *LH* benchmark, a total of 2, 669, 328 docking experiments were required to compute the IFPPs of all molecules in the benchmark. This included 144 pairs of ligands and their 499 decoys docked in 37 pockets. In real applications, solving a given scaffold hopping problem by screening 100.000 molecules (i.e. medium-size chemical libraries) would require 3, 700, 000 docking experiments, which is accessible. Nevertheless, screening of very large chemical libraries (millions of molecules) using IFPP molecular representations would lead to computational burdens. Therefore, in the following section, we introduce an alternative method based on the concept that isofunctional molecules tend to cluster together in the IFPP space, despite being distant in a chemical structure-defined space. This approach circumvents the computational constraints associated with computing the IFPP directly, facilitating the pre-screening of very large compound libraries.

### 2.3 Predicting IFPP similarity through Deep Metric Learning

We proposed to solve scaffold hopping problems by searching for molecules with similar IFPPs to that of a reference hit molecule, which requires to compute these IFPPs. One idea to bypass this calculation would be to train a machine learning algorithm that predicts the similarity of molecules in the space of IFPPs, without calculation of the IFPPs themselves. This approach uses concepts introduced in the domain of Metric Learning.

The principle of Metric Learning is to approximate a real-valued metric through an algorithm trained on available examples. This strategy has been applied in many fields, from face recognition ^23^ to representation learning. ^24^ Recently, Deep Neural Networks (DNN) have been employed for this purpose: they are trained to learn an embedding space in which the distance between points mimics the real-valued metric in the original space.

In our case, starting from the 2D structure of molecules as input, a Metric Learning approach would learn a new representation of the molecules such that, in this abstract space, the similarity between two molecules matches that of their similarity in Interaction Fingerprint Profiles. The learned embedding space serves as a surrogate of the IFPP space and can be interpreted as a dimension reduction of the IFPP.

#### 2.3.1 Siamese Networks

We chose a simple architecture that relies on the same principle as Siamese Networks. ^25,26^ This architecture consists in a network that takes distinct inputs, here two molecules, and outputs representations for each input, for which similarity or distance metrics can be computed. Figure 4 displays the global architecture that was adopted.

**Figure 4:**
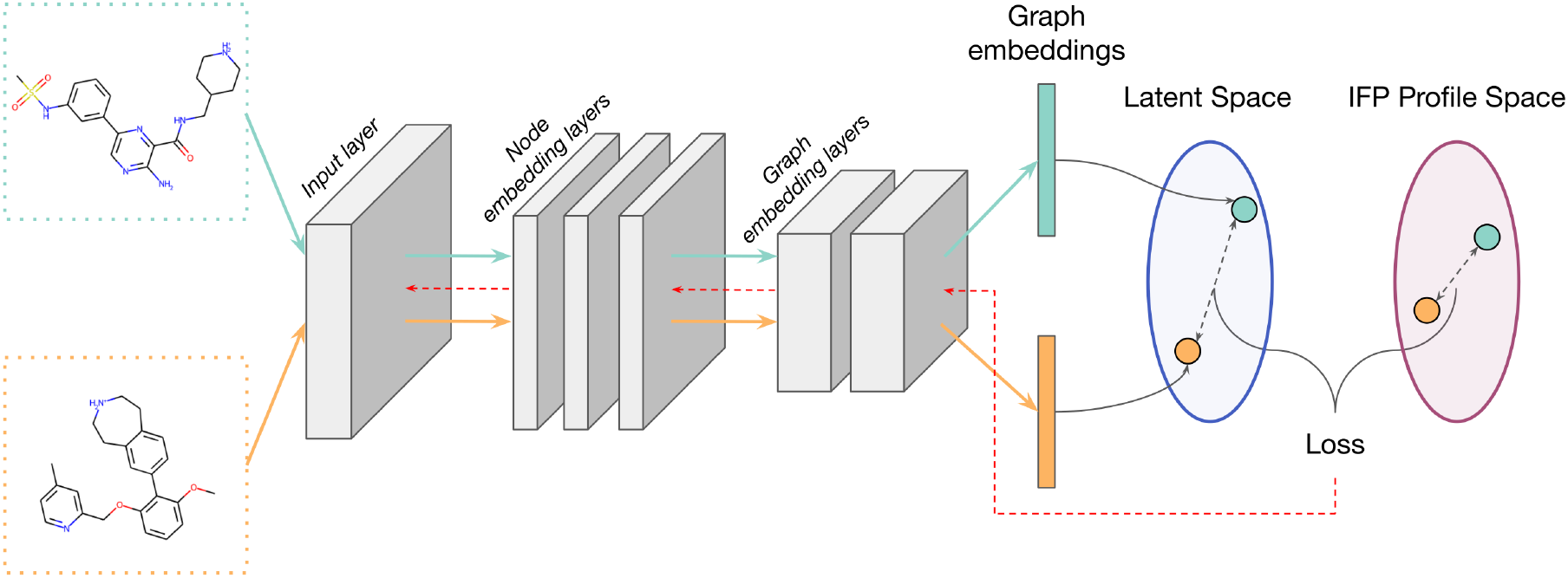
Illustration of the network architecture. A Siamese Neural Network is used to get the graph embeddings of pairs of molecules using Attentive FP. The GNN is composed of an input layer, node embedding layers and graph embedding layers. The similarity between molecules in the latent space are compared to their similarity in the IFPP space to compute the loss and train the model, as illustrated by the red arrow.

#### 2.3.2 Graph Neural Network for molecule embedding

We used Attentive FP, a Graph Neural Network (GNN) using the attention mechanism proposed by ^27^ to encode molecules with a readout function to obtain a graph representation for each molecule. This architecture achieved state-of-the-art predictions to a wide range of molecular properties. ^27^ It takes into account edge embeddings in the message passing steps of the nodes while relying on an attention mechanism. To get the final graph embedding of a molecule, the entire graph is treated as a supervirtual node that goes through stacked attentive layers which output a state vector for the whole molecule. We provide additional details in the section 4.4.

#### 2.3.3 Training the Deep Learning model with a Loss function

The deep learning model is trained so that in the learned feature space, the similarity between embeddings of molecules is similar to that computed with the IFPP of molecules.

Given a set of *N* molecules of known IFPP, we can define 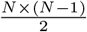 pairs, and compute their IFPP similarity, defined as the Tanimoto coefficient of their fingerprints, as above. In addition, for each pair, the deep learning model provides two graph embeddings (GE), for which we can also compute a similarity according to the following formula:

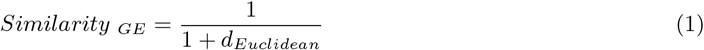

where *d*_*Euclidean*_ is the euclidean distance between the graph embeddings of a pair of molecules.

Training our deep learning model boils down to matching these two similarity measures. Therefore, we choose to compute the Root Mean Squared Log Error (RMSLE) between these two quantities, and use it as a part of the cost function to train our model:

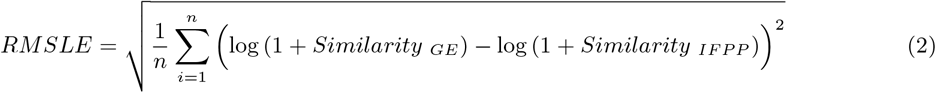

which can be rewritten as:

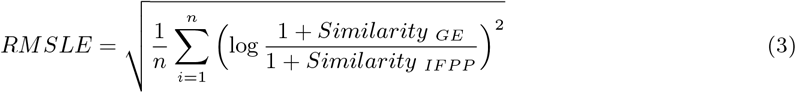

Where *n* is the batch size. This loss can be broadly interpreted as the relative error between the predicted and the actual similarities.

We also included the Kullback-Leibler divergence (*D*_*KL*_) to the RMSLE loss as a regularisation term to encourage the posterior distribution to be close to the prior distribution. ^28^ Therefore, the final loss used to train the model is defined as:

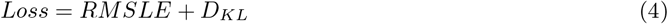

#### 2.3.4 Training Dataset

The model requires a training dataset of pairs of molecules with known IFPP.

In Section 2.2.3, we already computed the IFPPs for all actives and 144 *×* 499 = 71, 856 decoys in the *LH* benchmark. From the 71, 856 decoys, we randomly picked 60, 000 molecules to define the training set, and 10, 000 molecules to define the validation set, and the network was trained as detailed in Section 4.4

This allowed to train the model on pairs of molecules that are not considered with the *LH* benchmark. In this benchmark used to explore the efficacy of molecular representations, decoys and unknown actives are ranked according to their similarity with the known actives. Therefore, a training dataset containing only pairs of decoys ensures that the model is trained without any information about the actives, i.e. without using any pair of molecules taken into account for ranking. This limits potential bias in performance evaluation. Although the model will have seen decoys pairs during training, it will be tested only on pairs that include an unseen active molecule.

Once trained, this model can be employed to screen large compound libraries, allowing replacement of the expensive calculation of the IFPP by a quick inference of molecule embeddings, from which a predicted IFPP similarity is computed.

#### 2.3.5 Performance of the Metric Learning approach on *LH* Benchmark

The proposed deep learning model was evaluated on the *LH* benchmark to assess its ability to solve scaffold hopping cases. The protocol used to rank molecules is similar to that described in Section 2.2.3:

- for each scaffold hopping pair, one of the ligand is set as the known active and the other, the unknown active, is joined to 499 decoys
- the deep learning model is used to compute molecule embeddings
- according to this embedding, the similarities of decoys and unknown active with respect to the known active are computed
- decoys and the unknown active molecules are ranked according to their similarity with the known active
- the above steps are repeated by switching the active and unknown active roles to provide two scaffold hopping problems per pair of actives.

Table 2 gathers the success rate at 5% of the predicted IFPP similarity measure, in addition to those of other similarity measures considered above.

**Table 2:** Success rate at top 5% for similarity methods based on 3D descriptors and the Interaction Fingerprints Profile, and for the predicted Interaction Fingerprints Profile similarity.

| Method | Percent in Top 5% |
| --- | --- |
| 3D Shape | 14.9 |
| 3D Pharmacophore | 20.1 |
| IFPP | **24.7** |
| Predicted IFPP similarity | 19.8 |

As expected, the predicted IFPP similarity does not reach the performance of the similarity measure based on the true IFPP representation. We might argue that adjustments during the training phase and tuning of the model architecture may improve the performances. Still, this relatively simple model displays performances that reach those of the 3D Pharmacophore, while being much faster. Indeed, the 3D pharmacophore descriptors require computation of conformers and alignment of pharmacophores, leading to heavy calculations that are hardly compatible with screening of very large compounds libraries. On the contrary, once trained, the proposed Metric Learning approach quickly infers IFPP similarities, allowing large scale virtual screening campaigns. When screening very large compound libraries (millions of molecules), this approach could be used as a fast pre-screening campaign, keeping the top best few percent ranked molecule. The top-scoring molecules (up to hundreds of thousands of molecules) could be screened based on the computed IFPP representation. Note that in real-case applications, this representation could be used as input of any ligand-based approach, and not only with the simple similarity measure used in the present study.

## 3 Discussion

The main contribution in this article was to propose a novel molecular representation dedicated to the scaffold hopping problem: the Interaction Fingerprints Profile (IFPP). This profile is a representation that intend to capture possible binding modes of molecules, based on docking experiments in a panel of 37 diverse proteins. The IFPP was computed using a single docking protocol. This protocol was not optimized for each of the 37 proteins. However, the successful re-docking of the crystallographic ligand present in the binding sites of these proteins (see 4.3) indicates that the docking protocol was adapted to these structures, if not optimized. Future studies could consider evaluating several docking scores or using different docking software. This may allow to define consensus IFPPs with increased relevance with respect to the scaffold hopping problem.

We showed that increasing the number of proteins in the panel improves the performances of the IFPP similarity method on the *LH* benchmark. This indicates that the larger and the more diverse the protein set, the closer in the bioactivity space the IFPP would bring molecules that are distant in 2D structure. This also confirms the initial intuition that the proposed IFPP representation could be beneficial for scaffold hopping problems.

Due to the cost of docking used to define the IFPP, as it stands, this encoding is not scalable to screen large molecular libraries (millions of molecules), although in the case where a large library would be screened recurrently, it would be feasible to calculate this embedding only once.

To overcome this limitation, we proposed to leverage a Metric Learning approach to predict the IFPP similarity to the known active, which allows pre-screening of large molecular databases at a much lower computational cost. In a second step, screening using the computed IFPPs could be performed on prefiltered, and thus reduced, databases.

The interest of the IFPP representation was assessed according to the performance of its corresponding similarity measure on the *LH* benchmark. However, we would like to point that the proposed IFPP molecular representation is meant to be used as input in more sophisticated ligand-based method. Indeed, in real-life applications, unlike in this benchmark, several known active and inactive molecules would usually be available for the target of interest. Encoding molecules with the IFPP would allow to train QSAR or machine learning algorithms dedicated to help solving scaffold hopping problems.

Interestingly, in the *LH* benchmark, the 3D structures of the proteins targeted by the 144 scaffold hopping pairs are known, because these pairs were initially extracted from protein-ligand complexes of known 3D structures available in PDBbind. ^20^ In such cases, docking would be the reference method to solve scaffold hopping problems. Therefore, we assessed the performance of docking on this benchmark. More precisely, for each pair of active, one molecule was considered as the known active, and the unknown active and the decoys were docked in the PDB structure of the known active, to reproduce a real-life screening situation with docking. Then, the unknown active and the decoys were ranked according to their docking scores. The success rate was defined as the percentage of cases were the unknown active was ranked in the top 5%. More details about the protocol are given in 4.3. Docking retrieves the unknown active in the top 5% in 28.9% out of 135 considered scaffold hopping experiments. This performance would probably be higher in real-life cases, when a single protein target is considered. The docking protocol would then be finely tuned for this particular protein, which was not done in our benchmark application. Assuming that they would still remain in the same range, these performances are modest. This illustrates that large-step scaffold hopping is on average a difficult task, even when the structure of the protein is known. Note that on the same subset of 135 scaffold hopping experiments, the IFPP similarity measure had a success rate 30.4%, which is higher, although comparable to docking.

However, we consider that the Interaction Fingerprints Profile is essentially a new string to the bow of available methods for addressing these challenging problems. This representation should not replace others, but should rather be used in conjunction with other existing methods. Indeed, as shown in the Results, combining IFPP and 3D pharmacophore similarities increases the success rate on the *LH* benchmark.

## 4 Materials and Methods

### 4.1 Choice of the panel of proteins to define the IFPP molecular representation

Ideally, we would like to dock in a large panel of diverse proteins as it would lead to the most complete sampling of possible binding modes for molecules. However, this is not realistic due to the cost of computing the resulting IFPP representation because:

- the proteins in the panel must be suitable for reliable docking experiments, which (among other criteria) requires careful preparation of its pocket and successful re-docking of the crystallographic ligand, to ensure that the docking protocol is adapted to this pocket.
- docking in too many pockets to compute the IFPP would be very costly.

Therefore, to define a diverse but limited set of proteins, we considered the PDBbind database ^20^ containing 19.443 PDB files (2020) of 3D crystallographic structures of protein-ligand complexes. Only structures with resolution below 2.5 Å, and in which a drug-like ligand was bound, were kept, in order to keep only proteins with binding sites that are suitable for docking experiments with drug-like molecules. We selected complexes in which the ligand satisfied the following physicochemical parameters:

- No atoms other than H, C, N, O, F, P, S, Cl, Br
- 400g/mol ≤ Molecular weight ≤ 900g/mol
- -7 ≤ logP ≤ 7
- Maximum ring size = 7
- No more than 7 rotatable bonds (to simplify docking and conformer sampling)
- TPSA ≥ 30
- QED ≥ 0.3

These conditions exceed the criteria for drug-likeness, but they help removing unwanted chemotypes, such as salts, solvent or other molecules present in crystallisation buffers, and large interacting partners like peptides. This led to 1, 248 PDB structures of complexes involving 378 different proteins.

These structures include pan-inhibitors or ligands belonging to the same chemical series. In order to avoid redundancies in terms of pockets, we clustered ligands according to their Morgan similarity with a threshold of 0.4. For each resulting cluster, we only keep the PDB with the best resolution. This led to 872 PDB structures for 358 different proteins. We retained only one structure per protein, keeping the PDB structure with the ligand of largest molecular weight, in order to define the largest binding site for the subsequent docking experiments, leading to 358 structures. For these structures, we re-docked the crystallographic ligands in their corresponding pockets, and only kept those for which the docking poses had a RMSD below 2Å with respect to the crystallographic, ensuring that the docking protocol was adapted to these pockets. The final set of proteins should also avoid strong bias towards a few extensively studied families of proteins such as transferases or hydrolases, constituting almost 70% of crystallised PDBs. ^29^ Therefore, to assess structural diversity of the proteins belonging to the final panel, we used the SCOP database. ^19^ This database provides a hierarchical classification of proteins according to their 3D fold. The proteins unrecognised by the SCOP database (using the UniProt ID) were discarded, leaving us with 283 different proteins. We observed that our set was still highly enriched in some protein structural families, such as some of the kinases folds. In order to form a panel of diverse proteins, we kept only one PDB structure per superfamily. A superfamily, as defined by the SCOP database, gathers proteins with different sequences, but that may have a common evolutionary origin according to their structures and functional features. This filtering process left 73 superfamilies (so 73 PDB structures) defined in the SCOP database. To evaluate the effectiveness of the IFPP using the *LH* benchmark, we excluded 4 proteins that were also included in the benchmark to prevent bias. Overall, these successive filters led to 69 PDB structures of proteins that belong to various superfamilies. Information on this panel of proteins can be found in Supplementary Material.

### 4.2 Detecting protein-ligand interactions to compute the IFPP

The IFPP of a molecule is computed based on the interactions detected when docking this molecule in the panel of proteins. These interactions are identified using an in-house algorithm. Details about the type of interactions as well as their retained detection thresholds are provided in the Materials and Methods of our previous article. ^15^ Briefly, we used PLIP, ^30^ a publicly available software to detect interactions, including hydrogen bond, weak hydrogen bond, halogen bond, salt bridge, hydrophobic, pi-cation, and pi-stacking. Additionally, we added some interactions that are often missed: pi-hydrophobic, pi-amide, and multipolar. The code used to detect these interactions is available at: https://github.com/iktos/structure-interactions. For each protein of the panel, the interaction fingerprint associated is of fixed size for all molecules. Specifically, it corresponds to the number of residues in the binding site times the number of possible interactions (10). The binding site is defined as all residues within a radius of 10 Å from the crystallographic ligand of the protein. The final IFPP is obtained by concatenating the fingerprint determined for each protein of the panel.

### 4.3 Docking Protocols

In the manuscript, docking was used in two different contexts: as a tool used to compute the IFPP representation of molecules, and as a method assessed to solve scaffold hopping problems in the *LH* benchmark. The protocols employed in these two applications are described in the following.

#### Docking in the panel of proteins to compute molecular IFPPs

For all considered PDB structures, the ligand and the protein were prepared separately for docking with the DOCK6 software. ^18^ To prepare a protein-ligand structure, we applied the following steps:

- To simplify, all water molecules are removed.
- The molecule and the protein are protonated using the softwares SimulationPlus and PDB2PQR ^31^ respectively.
- We parametrise the complex for the molecular dynamics engine using Gromacs ^32^ with an Amber force field. ^33^
- We minimise the complex in vacuo so that all the hydrogens are well positioned.

When docking a molecule in a protein pocket with DOCK6, ^18^ only the pose with the best Grid-Based Score is retained, which relies on the non-bonded terms of the molecular mechanic force field.

This protocol was followed to select the panel 69 protein structures among those present in the PDBbing database (see above). The crystallographic ligand was re-docked in its pocket, and structures for which the RMSD between the re-docked ligand and the crystallographic structure was below 2.5 Å were kept. The same scoring function was also used to dock molecules in this protein panel, to ensures that the docking score was adapted to predict the best poses of molecules in the protein of this panel, and compute the resulting IFPP molecular representations.

This protocol was followed to prepare PDBs of the protein panel, and to define the IFPP of all molecules from the *LH* benchmark.

#### Protocol used to evaluate the performance of docking on the *LH* benchmark

The same protocol was used to prepare the PDBs corresponding to the pairs of active molecules in the *LH* benchmark, as to prepare those of the protein panel (see details above). In fact, for each scaffold hopping pair in the benchmark, two PDBs for the same protein are available, i.e. one complex for each of the two ligands. This preparation step did not succeed for all PDBs, and fixing the preparation steps in a case-dependant manner would have been extremely time-consuming. Thus, docking was assessed only for the 135 scaffold hopping experiments of the *LH* benchmark for which preparation of the PDBs succeeded. For each considered scaffold hopping case, the unknown active and the 499 decoys were ranked according to their docking score in the PDB structure of the known active. The performance of docking was assessed based on the percentage of cases for which the unknown active is ranked in the top 5%.

### 4.4 Architecture of the Metric Learning model

#### Model architecture

The architecture of the encoder used in the Metric Learning approach is based on Attentive FP. ^27^ First, a molecule is encoded by a 2D graph *G* = (*V, E*) where the nodes *V* represent the atoms and the edges *E* represent the bonds. Initial state vectors of length 128 for each node and edge are obtained with a fully connected input layer. Then, three GNN layers perform message passing on the node embeddings using an attention mechanism to include local information of the relevant neighbouring atoms. The message passing mechanism relies on a context vector incorporating neighbouring node and edge embeddings that goes through a gated recurrent unit GRU that updates the state vector of the node. To get the final embedding of the molecule, an initial molecule state vector is obtained by summing all node embeddings. Then, a readout block consisting in two attentive pooling layers is applied. In each pooling layer, a context vector of the molecule is computed using an attention mechanism on all node embeddings, which goes through a GRU that updates the molecule embedding. Finally, we get a graph embedding of length 256. We used the python packages `dgl-life` ^34^ and `PyTorch` ^35^ to implement the corresponding architecture.

#### Training phase

From the 71, 856 decoys of the *LH* benchmark, we randomly picked 60, 000 molecules to define the training set, and 10, 000 molecule to define the validation set. Pairs of molecules are formed during the training phase in each batch. We chose 64 as the batch size, so for each batch there are 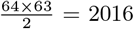 pairs. Each epoch consists in 937 batches, leading to 1, 890, 000 pairs formed at each epoch of the training. A learning rate of 0.0001 was used for *Adam* optimisation algorithm. We performed 200 epochs and kept the model with the lowest validation loss. Pairs are also formed inside batches for the validation, thus the model is evaluated on 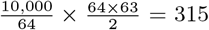 pairs. The size of graph embedding was set at 256 to encompass the intricacies of the IFPP.

