## supplementary for "A molecular representation to identify isofunctional molecules"

### Supplementary Material
### A molecular representation to identify isofunctional molecules

Philippe Pinel[1, 2, 3, 4], Gwenn Guichaoua[1, 2, 3], Nicolas Devaux[4], Yann Gaston-Mathé[4], Brice Hoffmann[4], and Véronique Stoven[1, 2, 3]

[1]Center for Computational Biology, Mines Paris-PSL, PSL Research University, Paris, France
[2]Institut Curie, Paris, France
[3]INSERM U900, Paris, France
[4]Iktos SA, Paris, France

| PDB | UniProt | Protein | Superfamily |
|---|---|---|---|
| **2qn1** | P00489 | glycogen phosphorylase | Type B glycosyltransferase-like |
| **3eq9** | P23687 | prolyl endopeptidase | Peptidase/esterase 'gauge' domain |
| **4bqh** | Q386Q8 | udp-n-acetylglucosamine pyrophosphorylase | Nucleotide-diphospho-sugar transferases |
| **3el8** | P00523 | tyrosine-protein kinase src | SH3-domain |
| **6ibx** | Q16875 | 6-phosphofructo-2-kinase/fructose-2,6-bisphosphatase 3 | Histidine phosphatase-like |
| **6i17** | Q16658 | fascin | Actin-crosslinking proteins |
| **6d55** | P01112 | gtpase hras | Ras-like P-loop GTPases |
| **4w9q** | Q13451 | peptidyl-prolyl cis-trans isomerase fkbp5 | TPR-like |
| **5dsx** | Q8TEK3 | histone-lysine n-methyltransferase, h3 lysine-79 specific | S-adenosyl-L-methionine-dependent methyltransferases |
| **3sff** | Q9BY41 | histone deacetylase 8 | Arginase/deacetylase-like |
| **4p5z** | P29320 | eph receptor a3 | galactose-binding domain-like |
| **1t48** | P18031 | protein-tyrosine phosphatase, non-receptor type 1 | (Phosphotyrosine protein) phosphatases II |
| **6duk** | P00533 | epidermal growth factor receptor | L domain-like |
| **6ccy** | P31749 | rac-alpha serine/threonine-protein kinase,piftide | PH domain-like |
| **3sfc** | P00797 | renin | Acid proteases |
| **6mob** | P10721 | mast/stem cell growth factor receptor kit | Protein kinase-like (PK-like) |

| | | | |
|---|---|---|---|
| **5bns** | P0A6R0 | 3-oxoacyl-[acyl-carrier-protein] synthase 3 | Thiolase-like |
| **4k9y** | Q05397 | focal adhesion kinase 1 | FAT domain of focal adhesion kinase |
| **6n0k** | O00625 | pirin | RmlC-like cupins |
| **5u8c** | P35439 | glutamate receptor ionotropic, nmda 1 | Type 2 solute binding protein-like |
| **5k13** | P10276 | retinoic acid receptor alpha | Nuclear receptor ligand-binding domain |
| **2ovx** | P14780 | matrix metalloproteinase-9 (mmp-9) | PGBD-like |
| **5syn** | O95372 | acyl-protein thioesterase 2 | alpha/beta-Hydrolases |
| **4k69** | P23946 | chymase | Trypsin-like serine proteases |
| **5nr7** | P9WKE1 | thymidylate kinase | P-loop nucleotide/nucleoside kinase-like |
| **6g2m** | Q9NPB1 | 5'(3')-deoxyribonucleotidase | HAD-like |
| **2ica** | P20701 | integrin alpha-l | vWA-like |
| **4pv5** | Q9CPU0 | lactoylglutathione lyase | Glyoxalase/Bleomycin resistance protein/Dihydroxybiphenyl dioxygenase |
| **4gvm** | P12497 | gag-pol polyprotein | N-terminal Zn binding domain of HIV integrase-like |
| **4ojr** | P03366 | hiv-1 integrase | DNA-binding domain of retroviral integrase |
| **6std** | P56221 | scytalone dehydratase | NTF2-like |
| **6qz8** | Q07820 | induced myeloid leukemia cell differentiation protein mcl | Bcl-2-like inhibitors of programmed cell death |
| **3dpe** | P22894 | neutrophil collagenase | Metalloproteases (zincins), catalytic domain |
| **5mw2** | P41182 | b-cell lymphoma 6 protein | POZ domain |
| **4ibj** | Q05127 | polymerase cofactor vp35 | Ebola VP35 IID-like |
| **5ufp** | Q99814 | endothelial pas domain-containing protein 1 | PYP-like sensor domain (PAS domain) |
| **6q9w** | O15151 | protein mdm4 | SWIB/MDM2 domain-like |
| 5t4b | P27487 | dipeptidyl peptidase 4 | DPP6 N-terminal domain-like |
| 4wp7 | P00352 | retinal dehydrogenase 1 | ALDH-like |
| 5ovg | Q07889 | son of sevenless homolog 1 | ENTH/VHS domain-like |
| 4yz9 | O75460 | serine/threonine-protein kinase/endoribonuclease ir | Ire1-RNaseL RNase domain-like |
| 6te6 | Q8TEK3 | histone-lysine n-methyltransferase, h3 lysine-79 specific | S-adenosyl-L-methionine-dependent methyltransferases |
| 5ur1 | P11362 | fibroblast growth factor receptor 1 | Immunoglobulin (Ig) domain-like |
| 6qed | P50579 | methionine aminopeptidase 2 | Creatinase/aminopeptidase catalytic domain-like |
| 5n9r | Q93009 | ubiquitin carboxyl-terminal hydrolase 7 | Cysteine proteinases |

| | | | |
|---|---|---|---|
| 3vhe | P35968 | vascular endothelial growth factor receptor 2 | Immunoglobulin (Ig) domain-like |
| 4l7n | Q9Y6F1 | poly [adp-ribose] polymerase 3 | WGR domain-like |
| 5twl | Q14680 | maternal embryonic leucine zipper kinase | UBA-like |
| 3hb4 | P14061 | estradiol 17-beta-dehydrogenase 1 | SDR-like |
| 6nss | P04629 | high affinity nerve growth factor receptor | Immunoglobulin (Ig) domain-like |
| 6f3i | Q9NWZ3 | interleukin-1 receptor-associated kinase 4 | DEATH domain |
| 2qcg | P11172 | uridine 5'-monophosphate synthase (ump synthase) | Ribulose-phoshate binding barrel |
| 2vwz | P54760 | ephrin type-b receptor 4 | SAM/Pointed domain |
| 6g2r | P08191 | type 1 fimbrin d-mannose specific adhesin | Bacterial adhesin-like |
| 4at4 | Q16620 | bdnf/nt-3 growth factors receptor | Immunoglobulin (Ig) domain-like |
| 3qrk | P00519 | tyrosine-protein kinase abl1 | alpha-Catenin/Vinculin-like |
| 4tsx | F2WR39 | integrase | Ribonuclease H-like |
| 4c9x | P36639 | 7,8-dihydro-8-oxoguanine triphosphatase | Nudix |
| 6qlr | P17931 | galectin-3 | Concanavalin A-like lectins/glucanases |
| 3tc5 | Q13526 | peptidyl-prolyl cis-trans isomerase nima-interactin | WW domain |
| 6as8 | Q1RBS0 | fml fimbrial adhesin fmld | Bacterial adhesin-like |
| 6r8w | P30405 | peptidyl-prolyl cis-trans isomerase f | Cyclophilin-like |
| 4kow | Q9HV14 | uncharacterized protein | Acyl-CoA N-acyltransferases (Nat) |
| 5d47 | P15090 | fatty acid-binding protein, adipocyte | Lipocalins |
| 5m6m | P98170 | e3 ubiquitin-protein ligase xiap | Inhibitor of apoptosis (IAP) repeat |
| 6q96 | Q00987 | e3 ubiquitin-protein ligase mdm2 | SWIB/MDM2 domain-like |
| 4z6i | Q9H8M2 | bromodomain-containing protein 9 | Bromodomain |
| 4lwu | P56273 | e3 ubiquitin-protein ligase mdm2 | SWIB/MDM2 domain-like |
| 1utr | P17559 | uteroglobin | Uteroglobin-like |

Supplementary Table 1: All 69 proteins used for the panel of proteins with the superfamily they belong to. The first 37, also in bold, constitute the panel for the IFPP evaluated on the whole *LH* Benchmark.

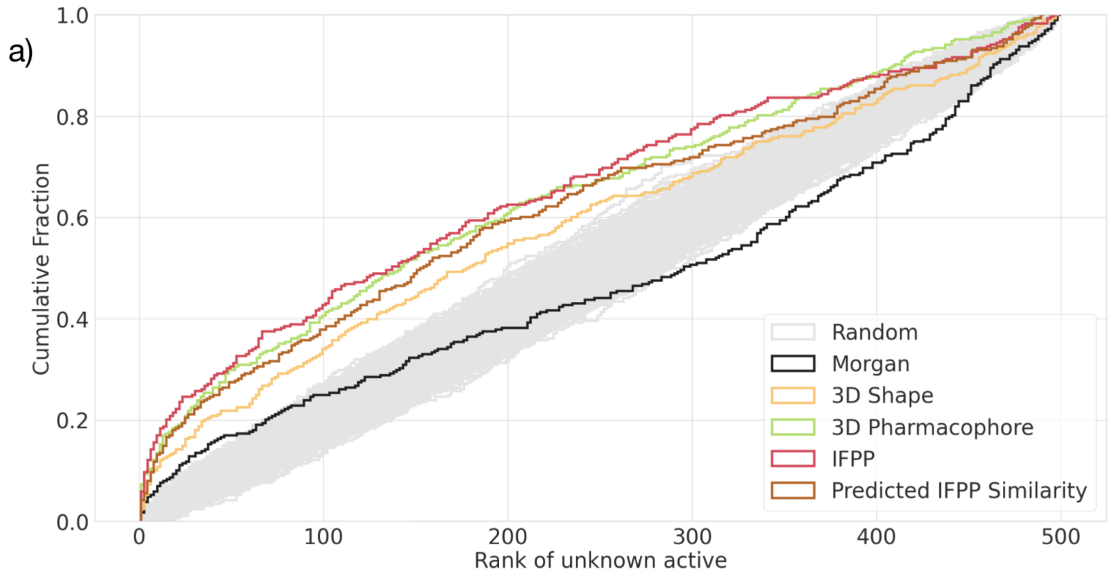


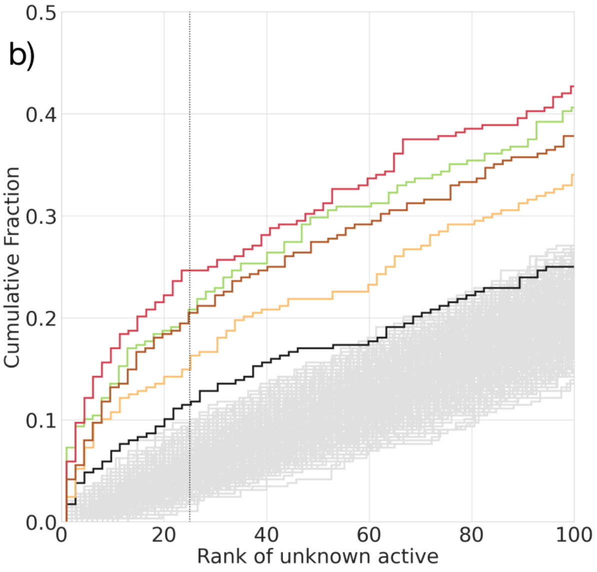


c)

| Method | Percent in Top 5% |
|---|---|
| Morgan | 11.5 |
| 3D Shape | 14.9 |
| 3D Pharmacophore | 20.1 |
| IFPP | **24.7** |
| Predicted IFPP Similarity | 19.8 |

Supplementary Figure 1: Results on the LH benchmark. The cumulative histogram curves of each similarity-based method are plotted in a). A zoom of the same graphs is provided in b). Table c) displays the percentage of successful scaffold hopping problems for similarity-based methods according to a rank of the unknown active in the top 5%.
